# The interplay between sulfur metabolism and desulfurization profile in *Rhodococcus*: Unraveling the role of the transsulfuration pathway

**DOI:** 10.1101/2022.02.14.480474

**Authors:** Olga Martzoukou, Panayiotis Glekas, Margaritis Avgeris, Diomi Mamma, Andreas Scorilas, Dimitris Kekos, Sotiris Amillis, Dimitris G. Hatzinikolaou

## Abstract

Biodesulfurization (BDS) is a process that selectively removes sulfur from dibenzothiophene and its derivatives. Several natural biocatalysts have been isolated, all harboring the highly conserved desulfurization operon *dszABC*. Even though the desulfurization phenotype is known to be significantly repressed by methionine, cysteine, and inorganic sulfate, the available information on the metabolic regulation of gene expression is still limited. In this study, scarless knockouts of the sulfur metabolism-related *cbs* and *metB* genes are constructed in the desulfurizing strain *Rhodococcus* sp. IGTS8. We provide sequence analyses for both enzymes of the reverse transsulfuration pathway and report their involvement in the sulfate- and methionine-dependent repression of the biodesulfurization phenotype, based on desulfurization assays in the presence of different sulfur sources. Additionally, the positive effect of *cbs* and *metB* gene deletions on *dsz* gene expression in the presence of both sulfate and methionine, but not cysteine, is uncovered and highlighted.

## Introduction

Microbial elimination of dibenzothiophene (DBT) and related organosulfur compounds, could allow for the biodesulfurization (BDS) of oil products by selectively removing sulfur from carbon-sulfur bonds, thus maintaining the calorific value of the fuel (1, 2). The process is mediated by the well-characterized 4S metabolic pathway that is found in several genera, with the most prominent that of *Rhodococci* (3). The three BDS genes are organized in a plasmid-borne operon, *dszABC*, and encode for a DBT-sulfone monooxygenase (*dszA*), a 2- hydroxybiphenyl-2-sulfinate (HBPS) desulfinase (*dszB*), and a DBT monooxygenase (*dszC*), respectively. A fourth chromosomal gene, designated *dszD*, encodes for an NADH-FMN reductase that energetically supports the pathway. One of the major disadvantages in exploiting the biotechnological potential of the BDS process is the sulfate, methionine, and cysteine-mediated transcriptional repression of *dsz* genes through a putative repressor-binding site in the *P*_dsz_ promoter. The operon is de-repressed in the presence of organosulfurs such as DBT and dimethyl sulfoxide (DMSO), and Dsz enzymes are considered sulfate-starvation-induced (SSI) proteins (4–6). However, information on the repression mechanism is still limited, and until recently, the sulfur assimilation pathways of *Rhodococci* had only been investigated *in silico* (7, 8). An exception is a very recent report that conducted comparative genomics and untargeted metabolomics analyses in *Rhodococcus qingshengii* IGTS8 and proposed a working model for assimilatory sulfur metabolism reprogramming in the presence of DBT (4).

### The Methionine-Cysteine interconversion pathways in bacteria

L-methionine and L-cysteine, the sulfur-containing amino acids responsible for *dsz* repression, are interconverted with the intermediary formation of L-homocysteine and L-cystathionine through the transsulfuration metabolic pathway.

L-methionine can be converted to L-homocysteine, an important intermediate of the transsulfuration pathway, via two possible routes (**Figure 1A**). The first requires the catalytic action of a methionine γ-lyase (MγL) for degradation to 2-oxobutanoate, NH_4_^+^, and methanethiol (9). The latter is then oxidized to sulfide, H_2_O_2_, and formaldehyde by a methyl mercaptan oxidase (MMO) present in *Rhodococcus* strain IGTS8 (10). A direct sulfhydrylation pathway can follow to convert sulfide to L-homocysteine, in condensation with either O-succinyl-L-homoserine (OSHS) or O-acetyl-L-homoserine (OAHS), through the catalytic action of MetZ (OSHS) or MetY (OAHS), thus serving as a precursor for sulfur-containing amino acid biosynthesis (4, 11). A second pathway for methionine catabolism, validated for Gram-positive bacteria, involves the sequential formation of S-Adenosyl-L-methionine (SAM), S-Adenosyl-L-homocysteine (SAH), and then L-homocysteine (4, 12–16). In the first step of the *forward* transsulfuration pathway, a γ-replacement reaction of L-cysteine and an activated L-homoserine ester (OSHS or OAHS) generates L- cystathionine and succinate or acetate, respectively, with the catalytic action of a Cystathionine γ-synthase (CγS; **Figure 1B**, Reactions M1 and M2) (17, 18). In the second forward transsulfuration step, L-cystathionine is acted upon by a Cystathionine beta-lyase (CβL) to form L-homocysteine and pyruvate.

**Figure 1.**
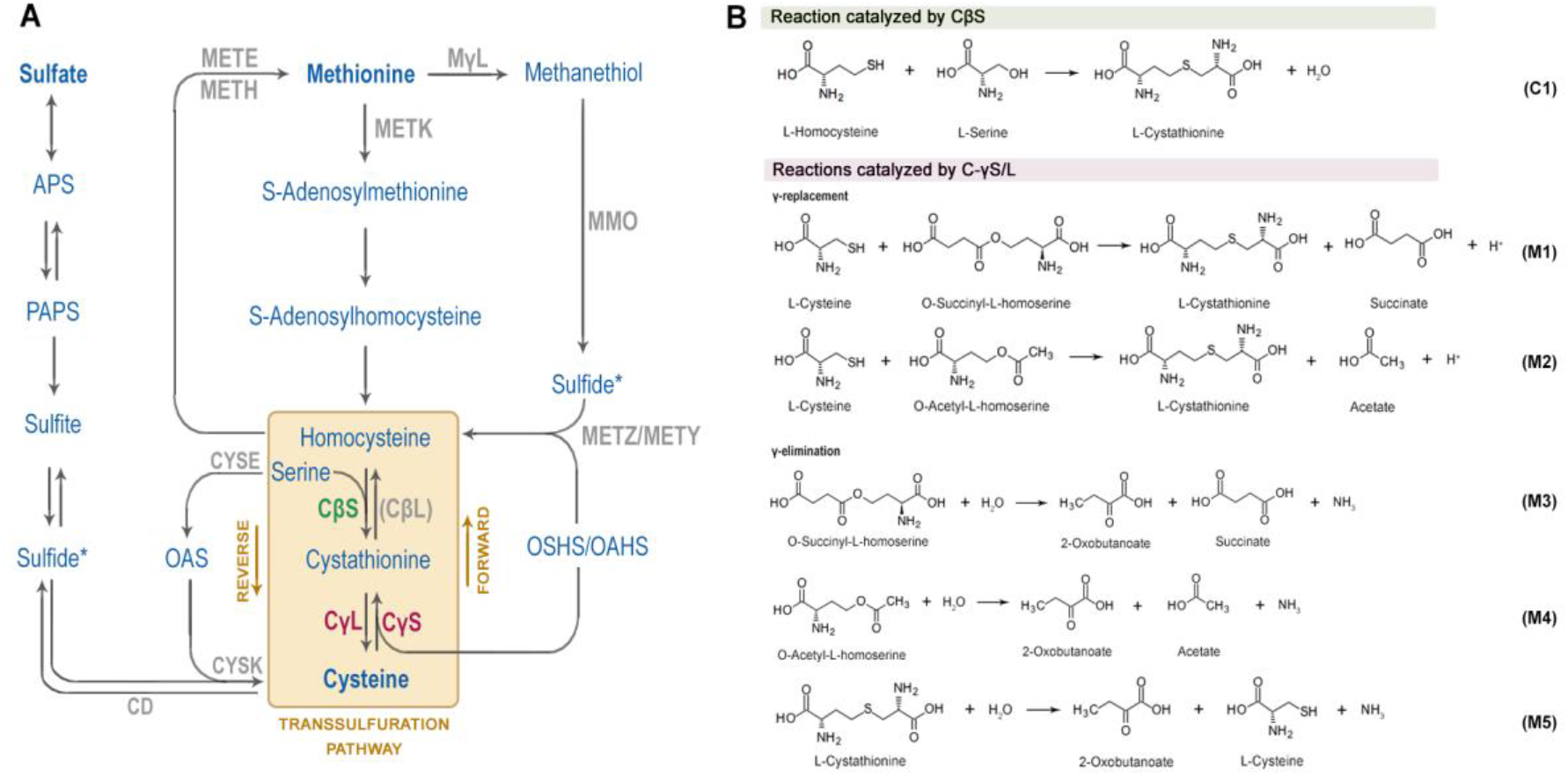
Bacterial sulfur metabolism. (A) Overview of Methionine and Cysteine biosynthesis and interconversion in bacteria as part of the sulfur assimilation pathway (APS: Adenylylsulfate, PAPS: 3’ Phosphoadenylyl sulfate, OAS: O-acetyl-L-serine, OSHS: O- succinyl-L-homoserine, OAHS: O-acetyl-L-homoserine). (B) Canonical reactions of sulfur metabolism catalyzed by CβS, METB (C-γS/L) in the *Corynebacteriales* order.

In Mycobacteria, L-homocysteine has been shown to be converted to L-methionine through a methylation step but, in general, serves as the precursor for L-cysteine biosynthesis via the *reverse* transsulfuration pathway (18). Therein, a Cystathionine β-synthase (CβS)- mediated condensation of L-homocysteine with L-serine generates L-cystathionine (**Figure 1B**, Reaction C1), which is then cleaved into L-cysteine, 2-oxobutanoate and ammonia, by a Cystathionine γ-lyase (CγL; **Figure 1B**, Reaction M5). Both key enzymes of the *reverse* transsulfuration pathway, CβS and, CγL, are pyridoxal phosphate (PLP)-dependent (19–22). This reverse transsulfuration metabolic route for L-Cysteine biosynthesis from L-Methionine has been also reported in mammals, yeasts, archaea and several other bacteria (13, 23–27). An alternative pathway for L-cysteine biosynthesis reported for *Corynebacterium glutamicum* and *Rhodococcus* strain IGTS8 requires the O-acetyl-L-serine (OAS) sulfhydrylase, CYSK, for the condensation of sulfide and OAS (4, 28). In the opposite direction, a reaction mediated by L-cysteine desulfhydrase (CD) leads to L-cysteine degradation into sulfide, pyruvate, and ammonia (28).

### Interconnection of transsulfuration and desulfurization pathways in Rhodococcus sp. and related species

The genome of the model desulfurizing bacterium, *R. qingshengii* IGTS8, harbors genes for CβS and C-γS/L, an indication for an active reverse transsulfuration pathway. The gene product of *cbs* gene is annotated as a putative CβS Rv1077, whereas the downstream located *metB* gene is predicted to encode a Cystathionine γ-synthase/lyase (C-γS/L). In one of the few reports providing information related to sulfur metabolism regulation in *Rhodococcus* sp., transposon mediated disruption of the *cbs* gene in the desulfurizing strain *R. erythropolis* KA2-5-1 led to induction of the *dsz* operon in the presence of sulfate and methionine, but not when cysteine or sulfite were used as sulfur sources (29). It was, therefore, suggested that sulfate and methionine are only indirectly involved in the repression of the *dsz* phenotype, contrastingly to cysteine and sulfite that are directly involved in the repression system. As inferred from the above referred literature, sulfur assimilation pathways and the regulation of *dsz* expression in response to different sulfur sources in desulfurizing *Rhodococcus* species, remains largely understudied *in vivo*.

Several genetic modifications were conducted with a direct biotechnological approach, aiming to increase the efficiency of BDS rather than elucidate the underlying sulfur assimilation regulatory mechanisms. As such, most of them engineer Gram-negative recombinant bacteria, such as *Escherichia coli* or *Pseudomonas* strains (30–32). However, a major limiting factor when *P. putida* CECT5279 was used as a biocatalyst in a biphasic system is the mass transfer rate of DBT from the oil to the aqueous phase (33). *P. putida* and all G(-) bacteria lack the robust hydrophobic cell wall of *Rhodococcus* and related mycobacterial species, and therefore DBT uptake from the oil phase is not efficient without the use of co-solvents (34, 35). This observation highlights the role of bacterial surface properties, such as hydrophobicity and cell wall thickness, for efficient BDS in biphasic media. In this regard, *Rhodococcus* biocatalysts that have the advantageous traits associated with the genus, pose as ideal candidates for genetic enhancement. However, this approach has not been favorable, especially in terms of targeted genetic modifications, owing to the extremely low amenability of *Rhodococcus* spp. to genome editing (36). To date, only a few studies have used genetically engineered desulfurizing *Rhodococcus* strains, which however harbor non-stable expression vectors or randomly integrated transposon elements (29, 37–40), while none have introduced site-directed, genome-based modifications in IGTS8 or in any other *Rhodococcus* sp. desulfurizing strain.

In the present work, we generate recombinant IGTS8 biocatalysts to investigate the effects of potential gene targets on biodesulfurization activity. More specifically, we implement a precise, scarless, two-step double crossover genetic engineering approach for the deletion of two sulfur metabolism-related genes, designated *cbs* and *metB*, located within the genome of *R. qingshengii* IGTS8 (41). Moreover, we provide sequence analyses of the related protein products (CβS and C-γS/L), with emphasis on highly conserved residues of the catalytic core, across various species. We present evidence that deletion of the *cbs* gene leads to the derepression of *dsz* phenotype mostly in the presence of sulfate, whereas the *metΒΔ* engineered strain seems to preferably desulfurize DBT when grown in the presence of methionine. Furthermore, we report the regulatory role of both CβS and METB (C-γS/L) in *dszABC* transcription levels in response to the presence of sulfate and methionine, but not cysteine.

Thus, we manage to indirectly mitigate the effect of sulfur source repression through targeted genome editing without modifying the native *dsz* operon.

## Results

### Sequence analysis of the cbs-metB genetic locus

In a previous study where random transposon insertion events were monitored in *R. erythropolis* KA2-5-1, it was shown that inactivation of *cbs* leads to high-level *dsz* genes expression in the presence of inorganic sulfate (29). However, under the same conditions, no transposon insertion events were isolated for the gene located downstream of the *cbs* locus, that of Cystathionine γ-synthase/lyase - C-γS/L (*metB*) (**Figure 2A**). Whole genome sequencing of *R. qingshengii* IGTS8 (41) revealed a 1386bp ORF for *cbs* and a 1173bp ORF for *metB*, with a similar organization to that of KA2-5-1 strain (29). The gene located upstream of the *cbs-metB* locus exhibits 61% identity with *M. tuberculosis* Rv1075c, a GDSL-Like esterase (46), while the gene downstream of *metB* is predicted to encode for an L-threonine ammonia-lyase. Analysis of the upstream flanking sequence of *cbs* suggests the presence of a bacterial promoter located ∼100 bp before the *cbs* start codon (**Figure 2B**).

**Figure 2.**
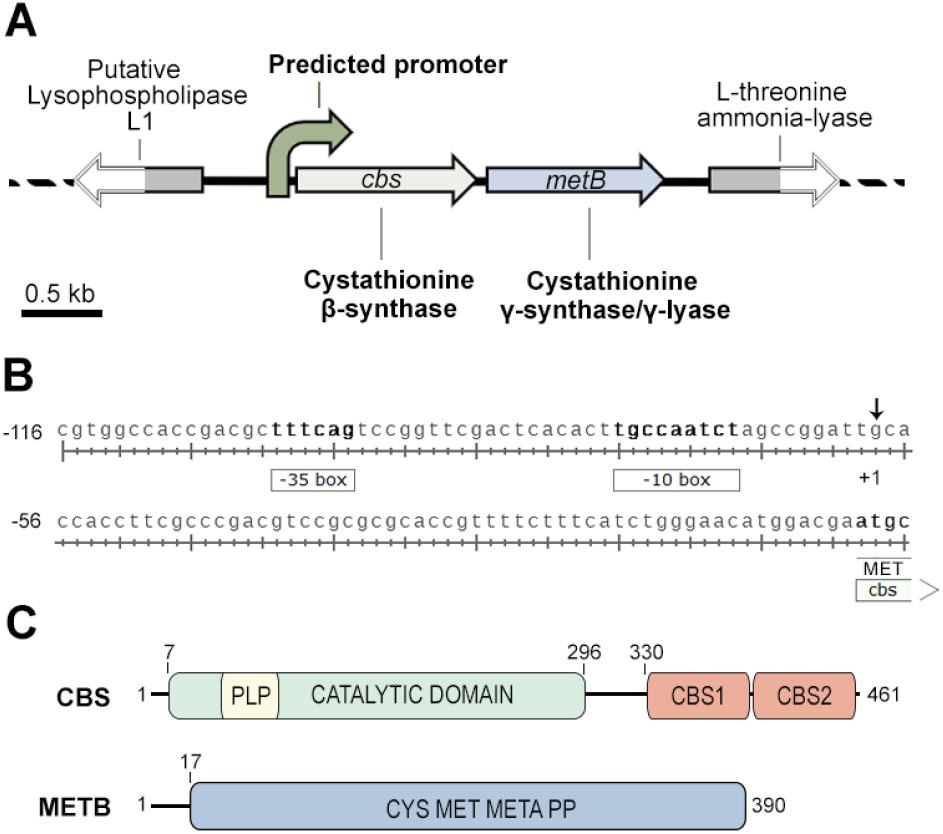
Properties of *cbs-metB* genetic loci and proteins. (A) Scheme of the *cbs-metB* gene cluster. (B) Bacterial promoter predicted sequence. -35 and -10 boxes are displayed, whereas an arrow indicates the predicted transcription initiation site (+1). (C) Schematic diagram of CβS and METB (C-γS/L) domain distribution. See main text for details.

Based on sequence homology, IGTS8 CβS consists of one N-terminal catalytic domain with the ability to bind PLP (7 - 296; pfam00291) and two C-terminal CBS regulatory motifs (CBS1, 330 - 397 and CBS2, 403 - 459; pfam00571) commonly referred to as the Bateman module (47, 48). In human and higher eukaryotes, the protein also harbors an N-terminal Heme binding domain of approximately 70 amino acid residues preceding the catalytic core domain, which has not been found in lower eukaryotes and prokaryotes (49–54). METB (C-γS/L) consists of a large Cysteine/Methionine metabolism-related PLP-binding domain (pfam01053), spanning almost the entire protein length (17–390) (**Figure 2C**). The translated amino acid sequences of IGTS8 CβS and METB were compared to other known CβS and C- γS/L proteins, respectively. Multiple sequence alignments revealed the presence of six conserved blocks in the catalytic core of CβS and three in the C-terminal Bateman module of the protein, whereas seven blocks are identified in METB (**Figure 3**). *M. tuberculosis* CβS shows the highest similarity score to IGTS8 CβS and shares extensive homology across the entire length of the protein (99% coverage, 83% Identity). Among the other known CβS homologs, MccA from *B. subtilis*, an O-acetylserine dependent CβS, shows a 41% overall identity for the compared region (65% coverage), although this protein completely lacks the C-terminal CBS1 and CBS2 regulatory domains. The *H. sapiens* and *S. cerevisiae* counterparts show 40% and 34% similarity, respectively, throughout both the Catalytic domain and the Bateman module of the CβS protein. Residues of the catalytic cavity that interact with CβS substrates and the cofactor PLP, are extremely well conserved across the compared sequences (**Figure 3A**, blue and yellow boxes respectively), whereas alignment of the C-terminal CβS regions reveals several highly conserved residues, distributed in three blocks (**Figure 3B**).

**Figure 3.**
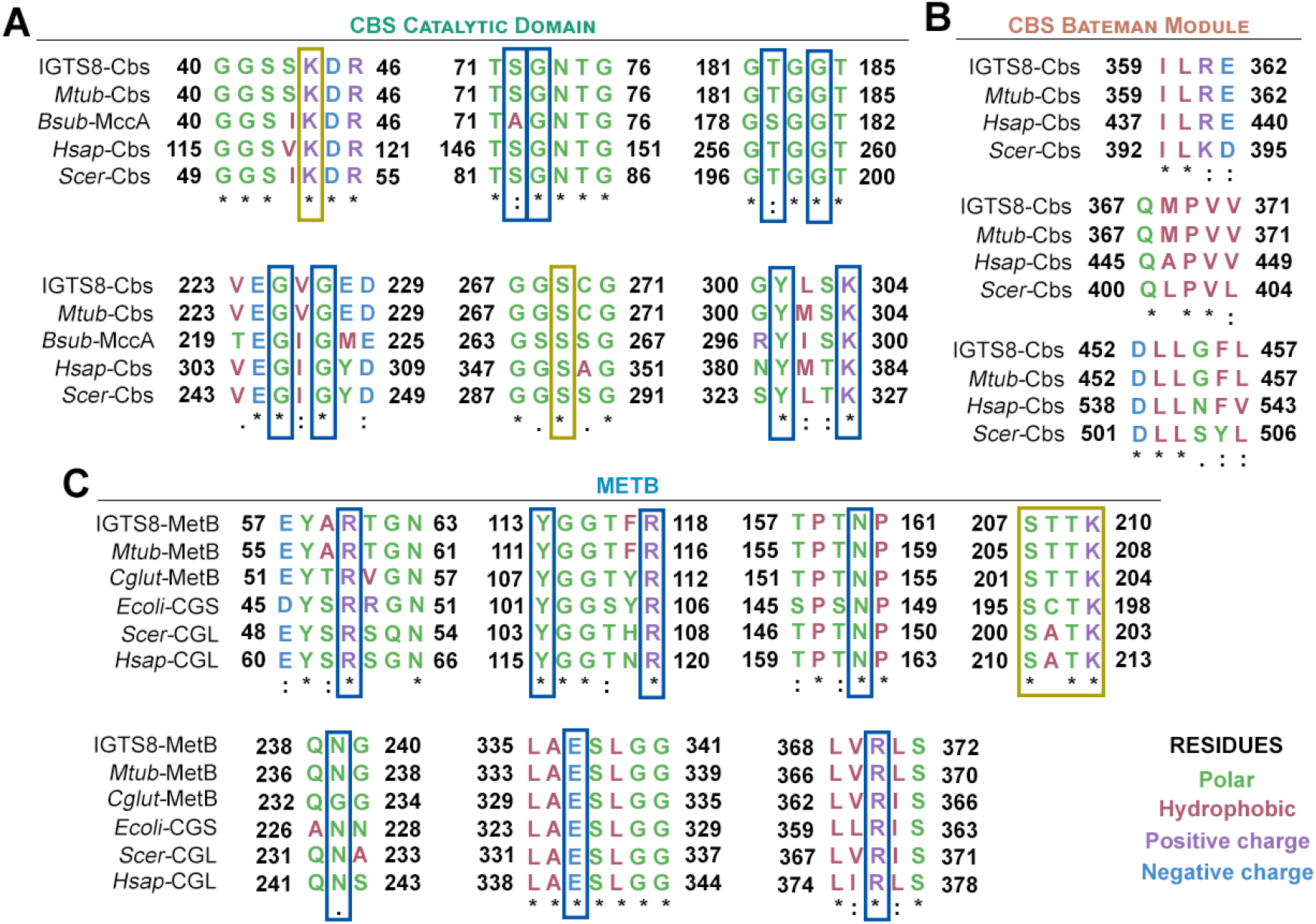
Multiple sequence alignments of CβS and C-γS/L, displaying only conserved residues configuring the active sites. (A) Comparison of *R. qingshengii* IGTS8 CβS with *M. tuberculosis* cystathionine β-synthase (Uniprot accession No: P9WP51); *B. subtilis* MccA (Uniprot accession No: O05393); Human CβS (Uniprot accession No: P35520-1); and *S. cerevisiae* CβS (Uniprot accession No: P32582). (B) Comparison of *R. qingshengii* IGTS8 METB with *M. tuberculosis* C-γS/L (Uniprot accession No: P9WGB7); *C. glutamicum* CγS (Uniprot accession No: Q79VD9); *E. coli* cystathionine γ-synthase (Uniprot accession No: P00935); *S. cerevisiae* cystathionine γ-lyase (Uniprot accession No: P31373) and Human cystathionine γ-lyase (Uniprot accession No: P32929). All multiple sequence alignments were done using ClustalO. Dashes indicate gaps introduced for alignment optimization. Asterisks (*) indicate fully conserved residues; double dots (:) denote strongly conserved residues and (.) show weakly conserved residues. Residues in yellow boxes participate in PLP-binding. Blue boxes denote residues involved in substrate binding (57, 22).

The METB (C-γS/L) multiple sequence alignment includes the *M. tuberculosis* and *C. glutamicum* METB, the cystathionine γ-synthase from *E. coli,* and the cystathionine γ-lyases from yeast and human (**Figure 3C**). *M. tuberculosis* and *C. glutamicum,* which are closely related to *R. qingshengii* IGTS8, possess homologs with the highest identity scores (73% and 65%, respectively), whereas coverage was high in all METB sequence alignments (95-99%). Notably, the CγS from the Gram-negative *E. coli* appears to have a lower similarity (42%) than the eukaryotic CγLs from *S. cerevisiae* and *H. sapiens* (49% and 47%, respectively). This observation is in line with the predicted bifunctionality of IGTS8 METB as both CγL and CγS, a unique feature that allows the synthesis of L-cysteine through L-methionine via the reverse transsulfuration pathway (18, 55, 56).

To study the role of CβS and METB (C-γS/L) in the regulation of *dsz* operon expression according to sulfur availability, scarless deletions of the corresponding genes (*cbs,* IGTS8_peg3012; and *metB,* IGTS8_peg3011) were performed with the use of the *pK18mobsacB* vector system (**Supplementary Figure S1A**; see also Materials and methods). Τhe isogenic knockout strains *cbsΔ* and *metBΔ* retained the ability to grow on liquid minimal media without supplemental cysteine or methionine, although the absence of CβS seems to have a negative effect on methionine-based growth (more details in the following paragraphs). We also tested the strains on solid minimal medium supplemented with 1 mM sulfate, DMSO, L-methionine, or L-cysteine. Our results indicate that all strains retain the ability to grow on basal salts medium (BSM), regardless of sulfur source addition (**Supplementary Figure S2B**). Thus, none of the constructed knockout strains is auxotrophic for methionine or cysteine, whereas none of the tested sulfur sources seems to lead to accumulation of intermediary toxic metabolites such as homocysteine, that could potentially inhibit growth (58).

### Ethanol is the preferred C source for maximum growth and desulfurization activity of IGTS8

To assess the effect of different carbon sources supplementation on BDS capability and determine the corresponding preferred carbon source for *R. qingshengii* IGTS8, we collected samples from actively growing cultures at three different time-points (early-log, mid-log, and late-log phase). Wild-type *Rhodococcus* cells were grown on either glucose, glycerol, or ethanol as sole carbon sources with 1 mM DMSO as sole sulfur source. Similarly to *R. erythropolis* KA2-5-1, the highest biomass and desulfurization activity for strain IGTS8 was obtained with the use of ethanol as a carbon source (59). On the contrary, utilization of glucose as a carbon source did not lead to a significant increase in biomass (0.12 ± 0.02 g/L) or to efficient BDS (0.30 ± 0.01 Units/mg DCW). In fact, cells did not exhibit a clear exponential growth even after 80 hours of incubation. The presence of glycerol as the sole carbon source led to a maximum biomass of 0.69± 0.06 g/L after 71 hours of growth and a BDS maximum of 19.00 ± 0.04 Units/mg DCW mid-log, still lower than the maximum biomass (0.88 ± 0.05 g/L) and catalytic activity (38.0 ± 1.9 Units/mg DCW) observed upon ethanol supplementation (**Figure 4**).

**Figure 4.**
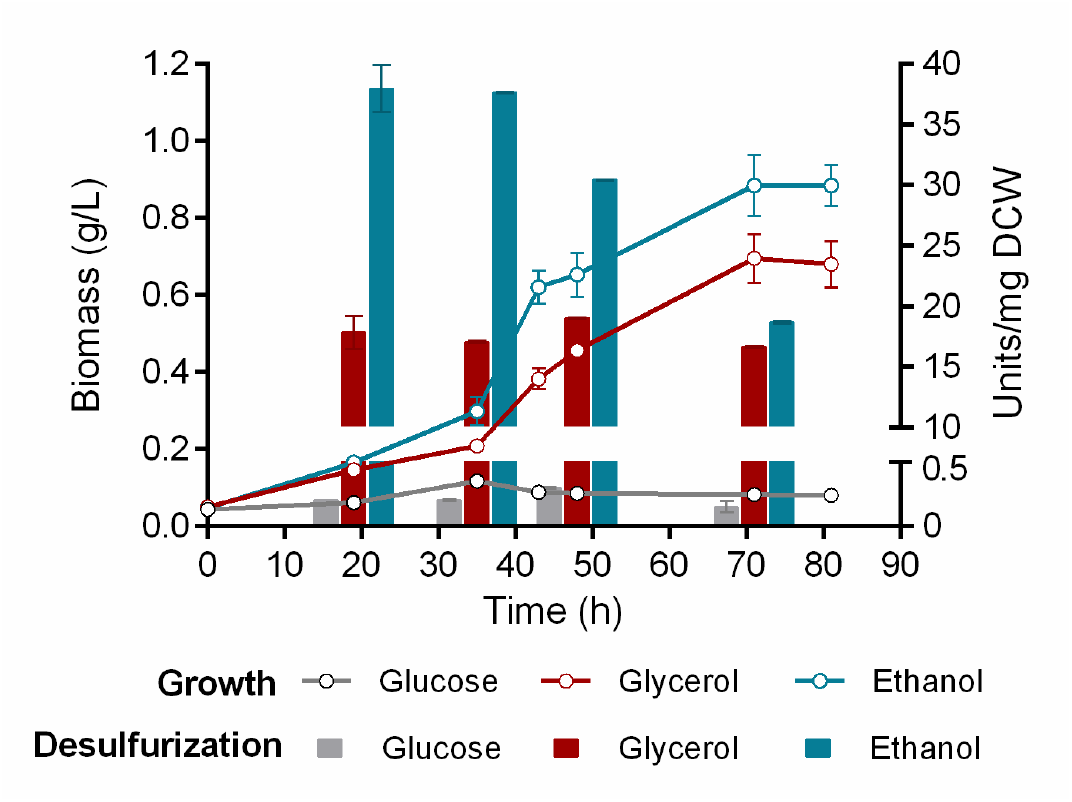
Ethanol is a preferred carbon source for *R. qingshengii* IGTS8. Effect of different carbon sources (0.055 M Glucose, 0.110 M Glycerol, 0.165 M Ethanol) on growth (Biomass; g/L) and BDS activity (Units 2-HBP/mg DCW) of *R. qingshengii* IGTS8. DMSO at a concentration of 1 mM was used as the sole sulfur source.

### Recombinant strains exhibit an altered desulfurization profile for repressive sulfur sources

We investigated the role of CβS and C-γS/L in desulfurization capability of *R. qingshengii* IGTS8, by comparison of growth rates and biodesulfurization (BDS) activity for isogenic *cbsΔ, metBΔ* and wild-type strains. All strains were grown in the presence of ethanol as the sole carbon source, whereas the BDS phenotype was firstly determined under non-repressive conditions, with the supplementation of DMSO at both low and high concentrations (0.1 and 1 mM, respectively; **Figure 5A-C**). For each separate strain, growth was not affected by the amount of DMSO added, indicating the limited S requirements for both wt and recombinant strains. The same can be contended for the determined specific BDS activity, which was also practically unaffected in each strain, by DMSO concentration. The growth yield of wt and *metBΔ* strains (0.91 ± 0.02 g/L and 1.25 ± 0.02 g/L, respectively; 0.1 mM DMSO) was increased compared to that of *cbsΔ* strain (0.70 ± 0.02 g/L; 0.1 mM DMSO). Comparison of respective maximum BDS activities (wt, 1mM DMSO: 29.0 ± 1.1 Units/mg DCW; *cbsΔ*, 0.1mM DMSO: 18.1 ± 1.9 Units/mg DCW; *metBΔ*, 0.1mM DMSO: 23.4 ± 2.0 Units/mg DCW) shows a possible negative effect of CβS depletion on growth and BDS, for cells grown on DMSO as the sole sulfur source.

**Figure 5.**
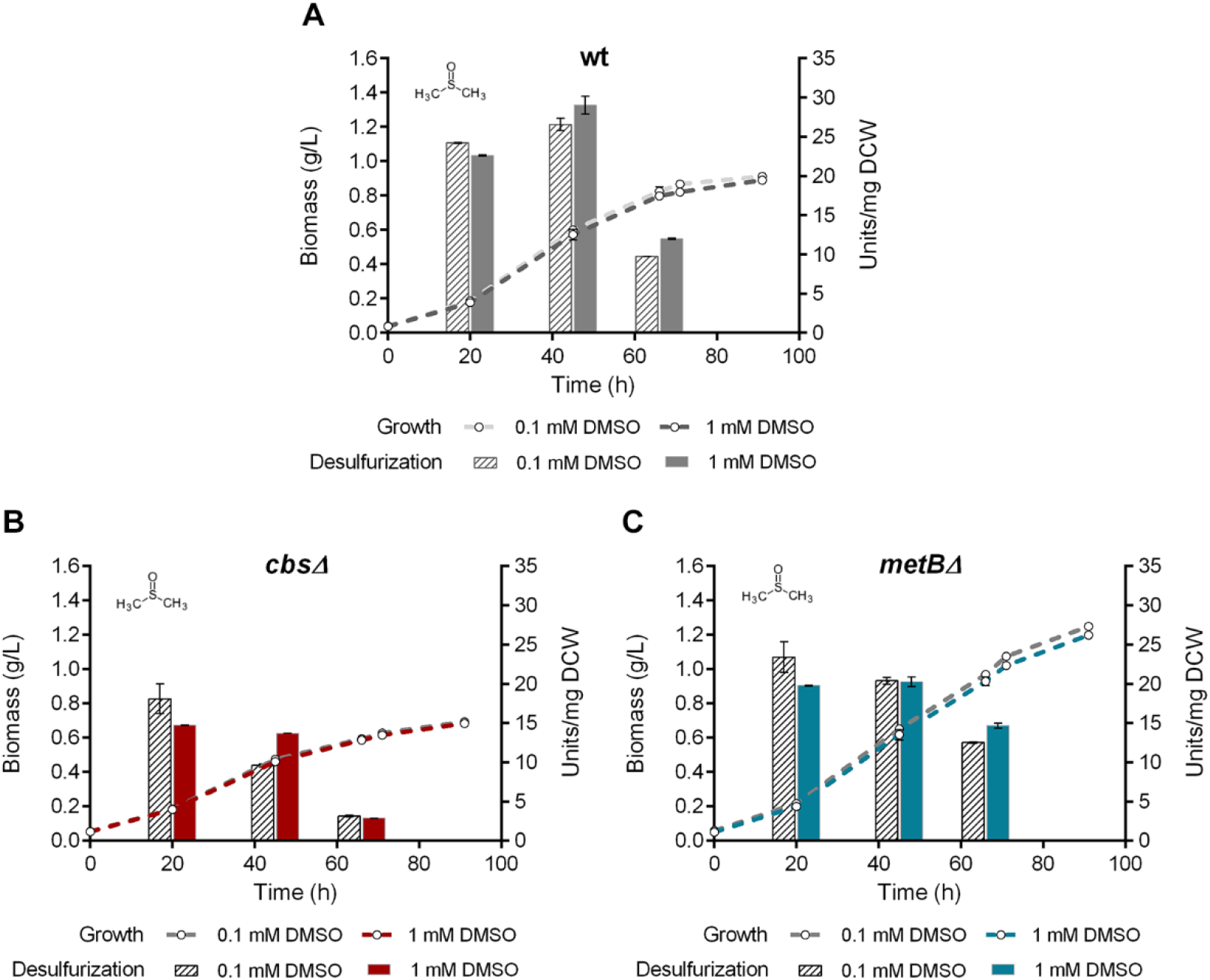
CβS and METB are not essential for growth and BDS in the presence of DMSO. (A-C) Growth (Biomass; g/L) and desulfurization capability (Units 2-HBP/mg DCW) of wt (A), *cbsΔ* (B), and (C) *metBΔ* strains, grown on CDM in the presence of low (0.1 mM) and high (1 mM) ***DMSO*** concentrations.

To determine the effect of *cbs* and *metB* gene deletions on cell growth and desulfurization activity in the presence of repressing sulfur sources, all strains were grown with the supplementation of sulfate, methionine, or cysteine as sole sulfur sources, at both low and high concentrations (0.1 mM and 1 mM). In all cases ethanol was used as the sole carbon source. Interestingly, the growth rate of each strain remains mostly unaffected by the concentration of the sulfur source, however, the maximum growth yield for all strains was achieved with the higher concentration of sulfate (1 mM; **Figure 6**) and with the lower concentration of the two sulfur-containing amino acids, methionine, and cysteine (0.1 mM; **Figure 7** and **Figure 8**, respectively).

**Figure 6.**
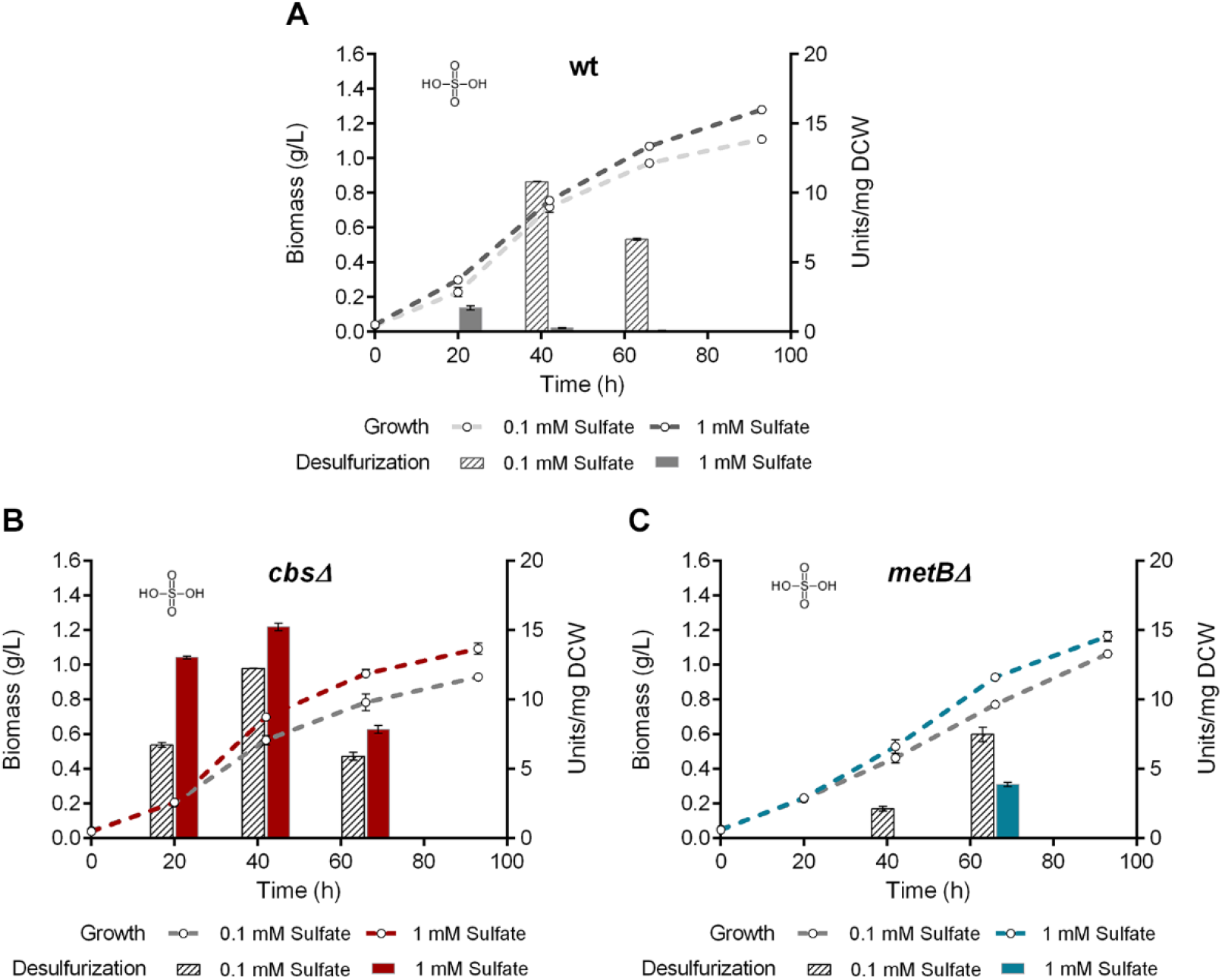
Recombinant strains desulfurize in the presence of sulfate. Growth curves (Biomass; g/L) and biodesulfurization efficiencies (Units 2-HBP/mg DCW) of wt (A), *cbsΔ* (B), and *metBΔ* (C) isogenic strains, in the presence of low and high ***sulfate*** concentrations as sole sulfur sources.

**Figure 7.**
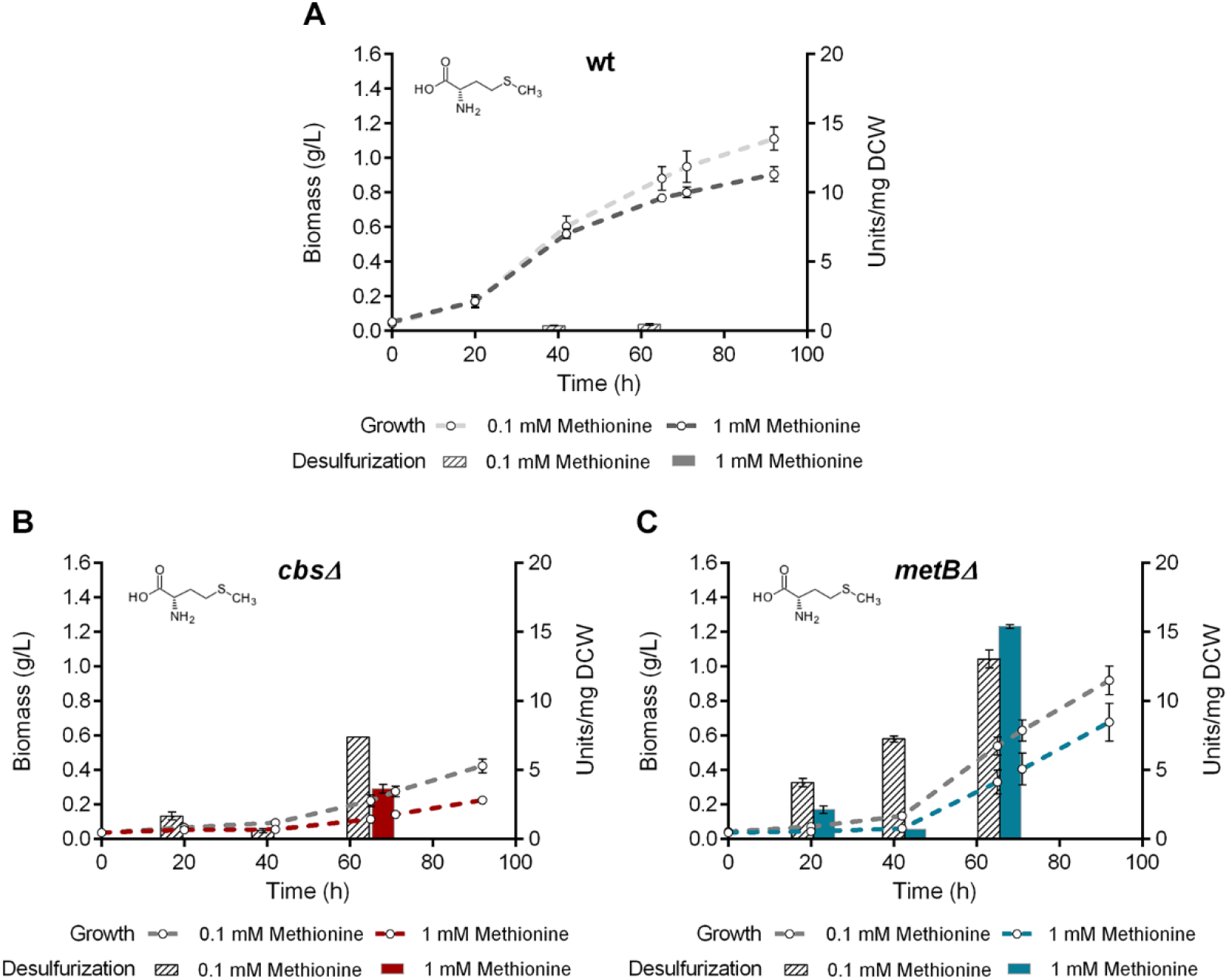
Methionine does not repress the BDS phenotype of recombinant strains. Growth curves (Biomass; g/L) and biodesulfurization efficiencies (Units 2-HBP/mg DCW) in the presence of low (0.1 mM) and high (1 mM) L-***methionine*** concentration, for wt (A), *cbsΔ* (B), and *metBΔ* (C) isogenic strains.

**Figure 8.**
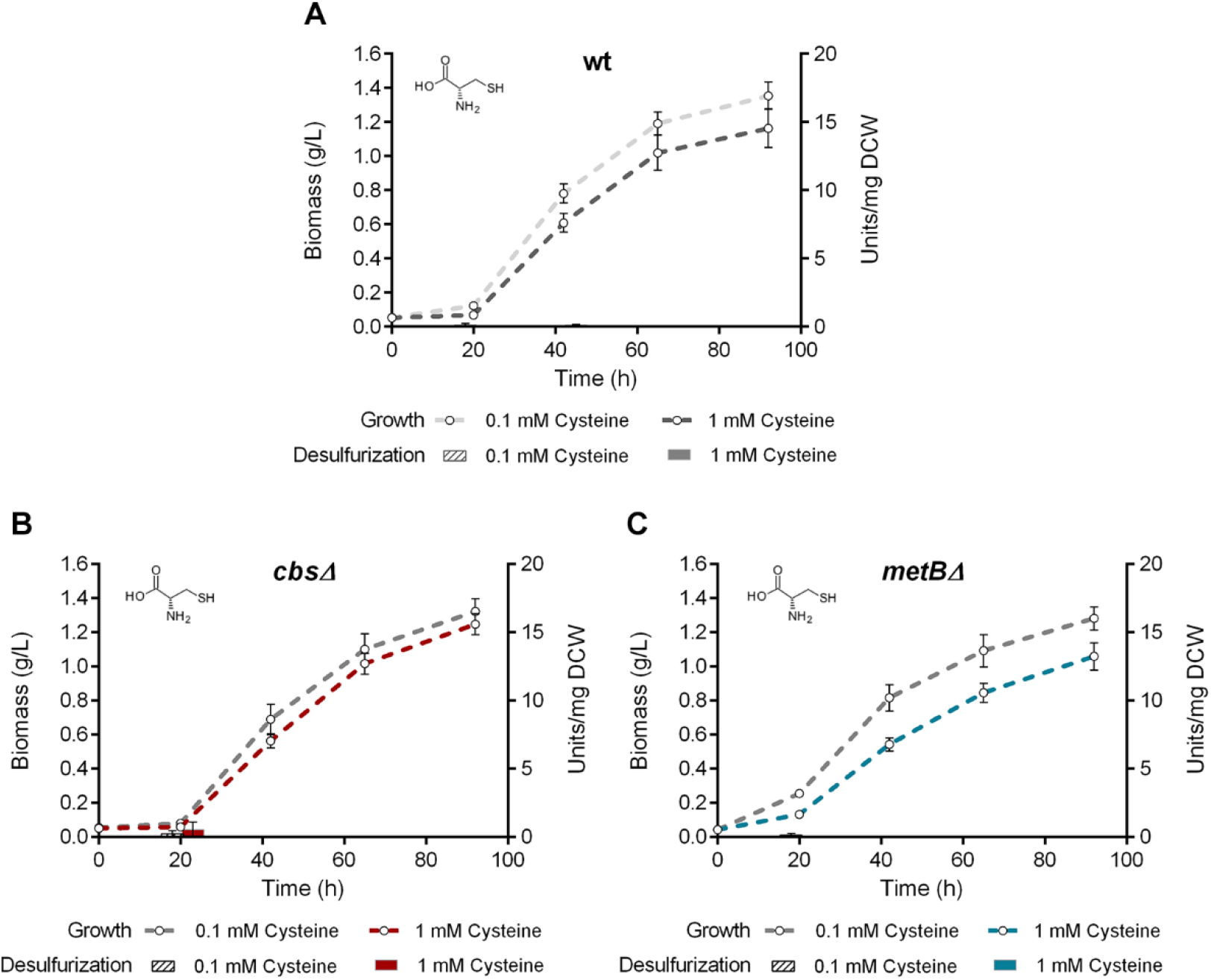
Cysteine negatively affects the biodesulfurization phenotype of wt and knockout strains. Growth curves (Biomass; g/L) and biodesulfurization efficiencies (Units 2-HBP/mg DCW) in the presence of low (0.1 mM) and high (1 mM) L-cysteine concentration, for wt (A), *cbsΔ* (B), and *metBΔ* (C) isogenic strains.

Sulfate addition in the bacterial culture efficiently represses the desulfurization phenotype of the wt strain, only when a high concentration is supplemented (1 mM; **Figure 6A**). Deletion of *cbs* leads to a slightly reduced biomass maximum compared to wt (1.09 ± 0.03 g/L and 1.28 ± 0.02 g/L, respectively), but enhances desulfurization 9-fold, reaching up to 15.23 ± 0.27 Units/mg DCW for *cbsΔ* in the presence of 1 mM sulfate, compared to 1.73 ± 0.17 Units/mg DCW for the wt strain (**Figure 6A and 6B**). The METB-depleted strain exhibits an intermediate maximum growth yield (1.17 ± 0.04 g/L), but a significant BDS phenotype is observed only during the late-exponential phase for 1 mM sulfate (7.49 ± 0.51 Units/mg DCW; **Figure 6C**).

One surprising finding is that the *cbsΔ* strain grows less efficiently than the wt and *metBΔ* strains, when methionine is used as the sole sulfur source (Maximum biomass for 0.1mM Methionine. wt: 1.11 ± 0.07 g/L; *cbsΔ*: 0.43 ± 0.04 g/L; *metBΔ*: 0.92 ± 0.08 g/L; **Figure 7**). In fact, this growth yield is one of the lowest observed for this strain. The BDS phenotype of the *cbsΔ* strain becomes evident after 65 hours of growth, while a preference for the low methionine concentration is also observed when comparing desulfurization activities (7.39 ± 0.01 Units/mg DCW for 0.1 mM, versus 3.65 ± 0.31 Units/mg DCW for 1 mM; **Figure 7B**). This observation is in line with the low growth yield reported for the transposon-disrupted *cbs* strain of *R. erythropolis* KA2-5-1, in the presence of 5 mM methionine (29). On the contrary, *metBΔ* strain can utilize methionine more efficiently than *cbsΔ* and exhibits important desulfurization activity after 65 hours of growth, with the addition of both low and high concentrations of the sulfur source (13.1 ± 0.65 and 15.4 ± 0.13 Units/mg DCW, respectively; **Figure 7C**). This phenotype is extremely interesting, especially when compared to the wt strain which is completely unable to desulfurize DBT, even in the presence of a low methionine concentration (0.46 ± 0.06 Units/mg DCW; **Figure 7A**).

Importantly, cysteine supplementation as the sole sulfur source in the culture medium results in complete inability of all three strains (wt, *cbsΔ* and *metBΔ*) to desulfurize DBT, even in the presence of low sulfur content (0.1 mM). However, growth is extremely efficient in all cases, reaching biomass maxima of 1.06 - 1.35 g/L after 90h of incubation. Based on this observation, it is highly likely that even low intracellular cysteine levels are an impeding factor in DBT biodesulfurization regardless of the reverse transsulfuration pathway functionality, possibly due to negative regulation of *dsz* operon expression (**Figure 8**).

### Gene deletion of cbs or metB leads to increased transcriptional levels of dszABC desulfurization genes in the presence of selected S sources

To elucidate the effect of *cbs* and *metB* deletions on the transcriptional levels of *dszABC* desulfurization genes, as well as the regulation of *dsz, cbs* and *metB* gene expression in response to sulfur availability, we performed a series of qPCR reactions for wt, *cbsΔ* and *metBΔ* strains under repressive and non-repressive conditions. In the presence of DMSO as sole sulfur source (**Figure 9A**), *dszABC* genes are efficiently expressed regardless of *cbs* or *metB* deletions. Additionally, *cbs* and *metB* transcriptional levels do not exhibit significant changes in the presence of DMSO. Sulfate or methionine supplementation (**Figure 9B** and **9C**, respectively) leads to repression of *dszABC* expression for the wt strain, while both *cbsΔ* and *metBΔ* knockout strains exhibit increased expression levels of the three desulfurization genes (*dszABC)*. Moreover, under the same conditions *metB* and *cbs* gene expression appears slightly elevated for the *cbsΔ* and *metBΔ* strains, respectively, compared to wt (**Figure 9B** and **9C**).

**Figure 9.**
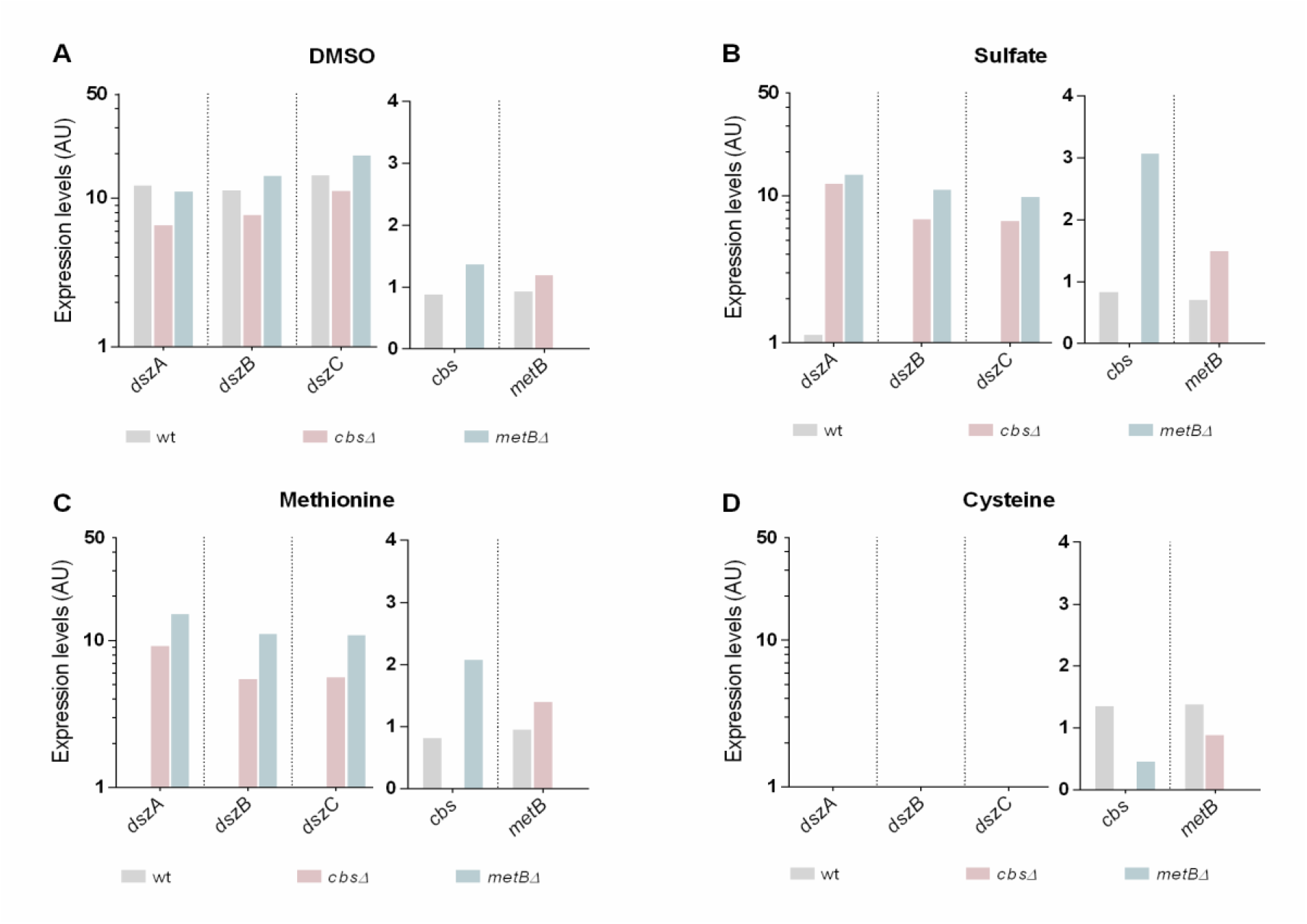
CβS and METB are critical for desulfurization genes *dszABC* expression in the presence of sulfate and methionine. Comparison of *dszA, dszB, dszC, cbs* and *metB* transcriptional levels for wt, *cbsΔ* and *metBΔ* isogenic strains, grown on 1 mM (A) DMSO, (B) Sulfate, (C) Methionine, or (D) Cysteine. Samples were collected from mid-log phase cultures (AU: Arbitrary Units; Relative expression levels compared to the calibrator sample. Logarithmic scale is used for *dszABC*. For details see Materials and Methods).

Interestingly, loss of *dszABC* transcription detected in the presence of cysteine (**Figure 9D**), not only for wt, but also for the two knockout strains. Furthermore, *cbs* and *metB* expression levels are slightly higher in the presence of cysteine for the wt strain, compared to other sulfur sources (**Figure 9A-C**). The results are in line with the observed sulfate- and methionine-related derepression of the desulfurization phenotype, in response to *cbs* and *metB* deletions.

## Discussion

Even though certain aspects of sulfur metabolism are generally well characterized in other Gram-positive bacteria, such as *B. subtilis,* the regulation of sulfur-assimilation-related gene expression remains unclear in *Rhodococcal* desulfurizing species. This is a curiously paradoxical situation, given that *R. qingshengii* IGTS8 is the most extensively studied biocatalyst for industrial biodesulfurization applications. The main reason for this is the challenging nature of genetic engineering for the actinomycete genus *Rhodococcus*, due to its high GC-content and prohibitively low homologous recombination efficiencies (36, 60). To our knowledge, no other studies have reported targeted, genome-based manipulations in desulfurizing *Rhodococci*. Contrastingly, plasmid-based modifications have been commonly used, which, however, are less preferred for industrial-scale applications as they exhibit a lower degree of genetic stability. Other approaches to date only include the *in silico* modeling of sulfur assimilation and the most recent proteomics and metabolomics analyses in strain IGTS8 (4, 8).

In the present work, we performed targeted and precise editing of the *R. qingshengii* IGTS8 genome for the first time, generating recombinant biocatalysts that harbor gene deletions of the two enzymes predicted to be involved in the reverse transsulfuration pathway. Importantly, primary amino acid sequence analyses of IGTS8 CβS and C-γS/L (METB), suggested the presence of highly conserved residue blocks that participate in active site configuration, binding of substrates and of the cofactor PLP. The high degree of similarity between the two IGTS8 enzymes and their respective counterparts found in the closely related species (61), *Mycobacterium tuberculosis* (83% Identity for CβS, 73% Identity for METB), suggests a conserved function for these two proteins as Cystathionine β-synthase and Cystathionine γ-lyase reverse transsulfuration enzymes, respectively. This result is in line with previous reports suggesting the existence of an operational reverse transsulfuration pathway for the genus *Rhodococcus* (4, 29).

In accordance with our sequence analyses and multiple alignment results, the sulfur assimilation model proposed by Hirschler et al. (4), also identified CβS as a cystathionine β- synthase and METB as a cystathionine γ-lyase. Interestingly, these authors also mention that sulfate addition in the culture medium leads to methionine production, and probably necessitates reverse transsulfuration metabolic reactions as the primary route for cysteine biosynthesis. Moreover, our study revealed a strategic role for CβS and METB in Dsz-mediated sulfur assimilation from organosulfates such as dibenzothiophene (DBT). The key role of CβS in sulfate- and methionine-mediated repression of the biodesulfurization phenotype agrees with previous results in a transposon-disrupted mutant of *R. erythropolis* KA2-5-1 (29). In the current study, however, the involvement of METB in the regulation of BDS is a novel finding. Our results indicate that biomass concentration and desulfurization capability are largely affected by the choice of sulfur source. This observation is in line with the findings of Hirschler et al. and Tanaka et al. (4, 29), as they report that the CysK-dependent alternative route for cysteine biosynthesis seems to be preferred under sulfate starvation conditions (BDS), however it is likely operating as a secondary pathway when sulfate is supplemented as the sole sulfur source. According to the same study, protein levels of CβS and METB were slightly higher, but not significantly different between the DBT and inorganic sulfate cultures. Based on our growth and desulfurization analyses, we further suggest that methionine supplementation resembles the sulfate-rich conditions and that the direct sulfhydrylation pathway is indeed operational in the background. This conjecture is based on the fact that during sulfate or methionine supplementation in the absence of CβS or METB, a complete cysteine deficiency would have manifested otherwise. However, knockout strain *cbsΔ* exhibits a preference for higher sulfate concentration, whereas methionine utilization appears to be less efficient. Overall, *cbs* deletion leads to a slower growth rate under all conditions tested, except for cysteine supplementation. In contrast, growth yield does not appear to be affected by the absence of METB, as evidenced by the growth rates observed for the *metBΔ* knockout strain.

Furthermore, *metB* deletion promotes BDS mainly in the presence of methionine and to a lesser extent, in sulfate-grown cells. This desulfurization phenotype follows the same pattern as the one observed for the strain harboring a *cbs* genetic deletion, in the presence of methionine as sole sulfur source. A possible explanation for this involves differential regulation of sulfur assimilation via actively operating alternative routes, given that *dsz* expression levels do not seem to differ significantly for either of the two knockout strains in the presence of sulfate or methionine. Τhe availability of sulfate is known to stimulate divergent routes for sulfate/sulfite reduction, while the latter serves as a metabolic branching point (4). Cysteine supplementation promotes growth for both knockout strains, which, however, do not exhibit the desulfurization phenotype. This is in line with the results reported by Tanaka et al. (29) for *R. erythropolis* KA2-5-1, as sulfate and methionine did not seem to be directly involved in the repression system, contrastingly to cysteine. Taking the suggested sulfur assimilation model into consideration, CβS and METB likely promote an increase of the free cysteine pool via the reverse transsulfuration pathway, when either sulfate or methionine is used as the sole sulfur source. This in turn could allow for *dszABC* efficient expression in *cbsΔ* and *metBΔ* strains, under sulfate- or methionine-rich conditions, given that sulfur-assimilation-genes expression is widely modulated in response to sulfur source availability (**Figure 10)** (4, 62).

**Figure 10.**
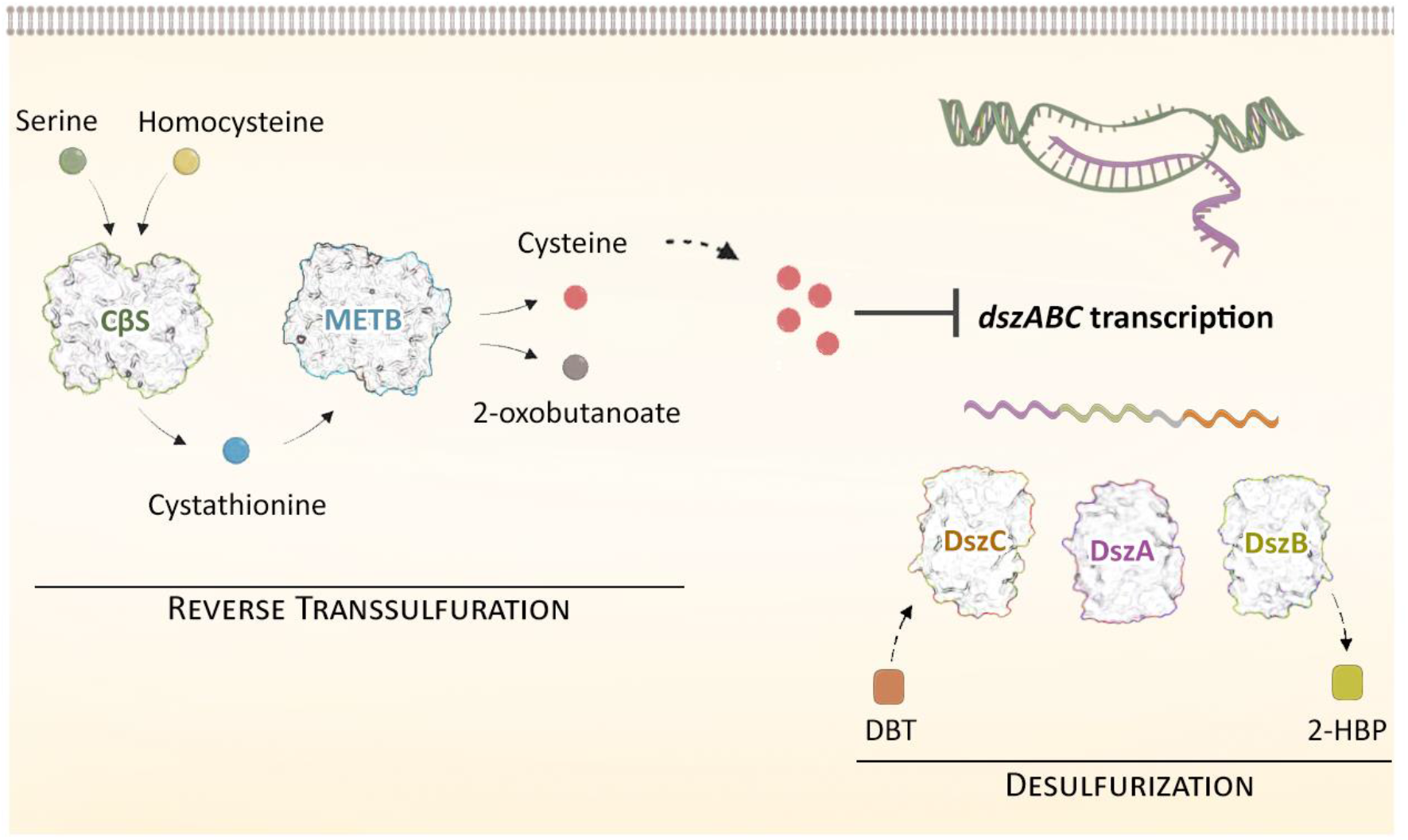
Proposed model illustrating the role of CβS and MetB (C-γS/L) in the regulation of desulfurization for *Rhodococcus qingshengii* IGTS8. Sulfate or methionine addition in the culture media, most likely necessitates reverse transsulfuration metabolic reactions as the primary route for cysteine biosynthesis. Fine-tuning of sulfur assimilation via intracellular cysteine levels is a common theme in bacterial species, where it seems to have evolved as a cellular mechanism to control gene expression appropriately, based on the available sulfur source type and abundancy. An increase in the free cysteine pool is suspected to exert an effect (directly or indirectly) on *dszABC* gene expression, leading to lack of biodesulfurization activity. Gene deletions of *cbs* or *metB*, abolish the cysteine-mediated *dsz* repression in the presence of selected sulfur sources, such as sulfate and methionine, thus leading to detectable transcript levels and biodesulfurization activity.

Based on these observations, the levels of *cbs, metB,* and *dszABC* genes expression were quantified in response to sulfur source supplementation. As evidenced by transcript level comparison for wt and *cbsΔ* or *metBΔ* knockout strains, CβS and MetB exert an effect on *dsz* gene expression, possibly via the regulation of cysteine biosynthesis through the reverse transsulfuration pathway. Deletions of the two genes might lead to reduction, but not depletion, of intracellular cysteine levels, promoting the expression of sulfur-starvation-induced proteins which in turn leads to the observed increase in biodesulfurization activity. This hypothesis is in line with the complete lack of biodesulfurization activity and the non-detectable *dsz* genes expression that was observed for wt as well as the knockout strains, in the presence of exogenously supplemented cysteine as the sole sulfur source.

Taken together, our approach focuses on the metabolic engineering of sulfur metabolism without manipulation of the 4S pathway genes. We thus propose the involvement of CβS and METB in the reverse transsulfuration pathway of *Rhodococcus qingshengii* IGTS8 and we validate the necessity of intact *cbs* and *metB* loci for the orchestration of *dsz-*mediated sulfur assimilation, in response to sulfur source availability.

## Materials and Methods

### Strains, growth conditions, and plasmids

The bacterial strains and plasmids used in this study are listed in Table 1. *Rhodococcus qingshengii* IGTS8 was obtained from ATCC (53968; Former names of the strain: *R. rhodochrous, R. erythropolis*). *Escherichia coli* DH5a and S17-1 strains were used for cloning and conjugation purposes, respectively. *Rhodococcus qingshengii* strains were routinely grown in Luria-Bertani Peptone (LBP) broth (1% w/v Bactopeptone, 0.5% w/v Yeast extract, and 1% w/v NaCl) at 30°C with shaking (180-200 rpm), or on LBP agar plates at 30°C. *E. coli* strains were grown in LB medium (1% w/v Bactotryptone, 0.5% w/v Yeast extract, and 1% w/v NaCl) at 37°C with shaking (180-200 rpm) or on LB agar plates at 37°C. Kanamycin (50 μg/ml) was used for plasmid selection in *E. coli.* Kanamycin (200 μg/mL) and Nalidixic acid (10 μg/ml) were used to select *R. qingshengii* transconjugants in the culture media. Counter-selection was performed on no-salt LBP (NSLBP) plates with 10% (w/v) sucrose.

**Table 1.** Bacterial strains and plasmids used in this study.

| Name | Description | Reference or Source |
| --- | --- | --- |
| **Strains** |  |  |
| *R. qingshengii* IGTS8 (wt) | DBT-degrading bacterium, Wild-type (wt) strain | ATCC 53968 |
| *cbs*Δ | Genetically engineered IGTS8 strain with *cbs* gene deletion | This study |
| *metB*Δ | Genetically engineered IGTS8 strain with *metB* gene deletion | This study |
| *E. coli* DH5α | F' Δ(lacZYA-argF) U169 *hsdR17* (rk<sup>-</sup> mk<sup>+</sup>) *recA1* *endA1* *relA1* | Laboratory stock |
| *E. coli* S17-1 | *recA* pro *hsdR* RP4-2-Tc::MuKm::Tn7 | ATCC 47055 |
| **Plasmids** |  |  |
| pK18mobsacB | Suicide vector derived from plasmid pK18; RP4 *mob*, *sacB*, KanR | Schafer et al., 1994 |
| pIGTS8*cbs* | Derived from pK18mobsacB for *cbs* gene deletion; RP4 *mob*, *sacB*, KanR | This study |
| pIGTS8*metB* | Derived from pK18mobsacB for *metB* gene deletion; RP4 *mob*, *sacB*, KanR | This study |

For growth tests on solid media, *R. qingshengii* cells were grown on basal salts medium (BSM) prepared according to (Karimi et al., 2017), containing 0.165 M ethanol (0.33 M carbon) as carbon source and 1% w/v agarose. Sulfur sources were supplemented at a final concentration of 1 mM S. For biodesulfurization studies, *R. qingshengii* wt and recombinant strains were grown on a sulfur-free chemically defined medium (CDM) containing 3.8 g NaH_2_PO_4_·H_2_O, 3.25 g Na_2_HPO_4_·7H_2_O, 0.8 g NH_4_Cl, 0.325 g MgCl_2_·6H_2_O, 0.03 g CaCl_2_·2H_2_O, 8.5 g NaCl, 0.5 g KCl, 1 mL Metal Solution, and 1 mL of Vitamin solution in 1 L of distilled water (pH 7.0). The metal solution was composed (per L of distilled water): Na_2_-EDTA, 5.2 g; FeCl_2_·4H_2_O, 3 mg; H_3_BO_3_, 30 mg; MnCl_2_·4H_2_O, 100 mg; CoCl_2_·6H_2_O, 190 mg; NiCl_2_·6H_2_O, 24 mg; CuCl_2_, 0.2 mg; ZnCl_2_, 0.5 mg; Na_2_MoO_4_·2H_2_O, 36 mg; Na_2_WO_4_·2H_2_O, 8 mg; and Na_2_SeO_3_·5H_2_O, 6 mg. The vitamin solution contained (per L of distilled water) calcium pantothenate, 50 mg; nicotinic acid, 100 mg; *p-*aminobenzoic acid, 40 mg; and pyridoxal hydrochloride, 150 mg. CDM was supplemented with sulfate, dimethyl sulfoxide (DMSO), L-methionine or, cysteine as the sole sulfur source (0.1 or 1 mM) and 0.165 M ethanol, 0.055 M glucose, or 0.110 M glycerol as carbon sources (0.33 M carbon), depending on the experiment. *pK18mobsacB* (Life Science Market, Europe) was used as a cloning and mobilization vector.

### Enzymes and chemicals

All restriction enzymes were purchased from TaKaRa Bio or Minotech (Lab Supplies Scientific SA, Hellas). Chemicals were purchased from Sigma-Aldrich (Kappa Lab SA, Hellas) and AppliChem (Bioline Scientific SA, Hellas). Conventional and high-fidelity PCR amplifications were performed using KAPA Taq DNA and Kapa HiFi polymerases, respectively (Kapa Biosystems, Roche Diagnostics, Hellas). All oligonucleotides were purchased from Eurofins Genomics (Vienna, Austria) and are listed in Table S2.

### Genetic manipulations and DNA sequence analysis

The genomic DNA of *Rhodococcus* strain IGTS8 was isolated using the NucleoSpin Tissue DNA Extraction kit (Macherey-Nagel, Lab Supplies Scientific SA, Hellas) according to the manufacturer’s instructions. Plasmid preparation and DNA gel extraction were performed using the Nucleospin Plasmid kit and the Nucleospin Extract II kit (Macherey-Nagel, Lab Supplies Scientific SA, Hellas). DNA sequences were determined by Eurofins-Genomics (Vienna, Austria). The online software BPROM was used for bacterial promoter prediction (http://www.softberry.com/cgi-bin/programs/gfindb/bprom.pl).

### Construction of knockout strains

Unmarked, precise gene deletions of Cystathionine β-synthase (*cbs*) or Cystathionine γ- lyase/synthase (*metB*) were created using a two-step allelic exchange protocol (43). Upstream and downstream flanking regions of the *cbs* gene of strain IGTS8 were amplified and cloned into the pK18mobsacB vector (44), using the primer pairs *cbsUp-F/Up-R* and *cbsDown- F/Down-R*, respectively, yielding plasmid pIGTS8*cbs*. Similarly, for the flanking regions of *metB* gene, primer pairs *metBUp-F/Up-R* and *metBDown-F/Down-R* were used to construct plasmid pIGTS8*metB*. *E. coli* S17-1 competent cells were transformed with each of the modified plasmids. *R. qingshengii* IGTS8 knockouts were created after conjugation (45) with *E. coli* S17-1 transformants, using a two-step homologous recombination (HR) process. Following the first crossover event, sucrose-sensitive and kanamycin-resistant IGTS8 transconjugants were grown in LB overnight with shaking (180 rpm), to induce the second HR event. Recombinant strains were grown on selective media containing 10% (w/v) sucrose and tested for kanamycin sensitivity, to remove incomplete crossover events. Gene deletions *cbsΔ* and *metBΔ* were identified with PCR and confirmed by DNA sequencing, using external primer pairs *cbs-5F-check-F/cbs-metB-3R-check* and *metB-5F-check/metB-3R-check*, respectively.

### Growth and desulfurization assays

For growth studies and resting-cells’ biodesulfurization assays, wild-type and recombinant *R. qingshengii* strains were grown in CDM under different carbon and sulfur source type and concentrations. Growth took place in 96-well cell culture plates (F-bottom; Greiner Bio-One, Fischer Scientific, US) with 200 μL working volume in thermostated plate-shakers at 30 °C and 600 rpm (PST-60HL, BioSan, Pegasus Analytical SA, Hellas). For each condition, an initial biomass concentration of 0.045-0.055 g/L was applied, while 20 identical well-cultures were used. Biomass concentration, expressed as Dry Cell Weight (DCW), was estimated by measurement of optical density at 600 nm with a Multiskan GO Microplate Spectrophotometer (Thermo Fisher Scientific, Waltham, MA USA), and calculations were based on an established calibration curve.

For the resting-cells biodesulfurization assays, the content of 2 to 4 identical well-cultures was harvested at early-, mid- and late-exponential phase, centrifuged at 3.000 rpm for 10 min, and the medium was discarded. Pellets were washed with a S-free buffer of pH 7.0 (Ringer’s), and cells were resuspended in 0.45 ml of 50 mM HEPES buffer, pH 8.0. Suspensions were separated into three equal volume aliquots (0.15 mL) in Eppendorf tubes. 0.15 mL of a 2 mM DBT solution in the same buffer were added in each tube, and desulfurization reaction took place under shaking (1200 rpm) for 30 min in a thermostated Eppendorf shaker (Thermo Shaker TS-100, BOECO, Germany). The reaction was terminated with the addition of equal volume (0.3 ml) acetonitrile (Labbox Export, Kappa Lab SA, Hellas) and vigorous vortexing. Suspensions were centrifuged (14.000xg; 10 min), and 2-HBP produced was determined in the collected supernatant through HPLC. One of the tubes, where the 0.3 mL acetonitrile was added immediately after DBT addition (t=0), was used as blank. Desulfurization capability was expressed as Units per mg dry cell weight, where 1 Unit corresponds to the release of 1 nmole of 2-HBP per hour. The linearity of the above-described assay with respect to biomass concentration has been verified for up to 2 h reaction time and up to 100 μΜ 2-HBP produced.

### HPLC analysis

High-performance liquid chromatography (HPLC) was used to quantify 2-HBP and DBT. The analysis was performed on an Agilent HPLC 1220 Infinity LC System, equipped with a fluorescence detector (FLD). A C18 reversed phase column (Poroshell 120 EC-C18, 4 μm, 4.6x150 mm, Agilent) was used for the separation. Elution profile (at 1.2 mL/min) consisted of 4 min isocratic elution with 60/40 (v/v) acetonitrile/H_2_O, followed by a 15 min linear gradient to 100% acetonitrile. Fluorescence detection was performed with excitation and emission wavelengths of 245 nm and 345 nm, respectively. Quantification was performed using appropriate calibration curves with the corresponding standards (linear range 10 - 1000 ng/mL).

### Extraction of total RNA

*R. qingshengii* IGTS8 wild-type, *cbsΔ,* and *metBΔ* deletion strains were grown in CDM medium containing DMSO, MgSO_4_, methionine, or cysteine as the sole sulfur source (1 mM S). Ethanol was used as a carbon source to a final concentration of 0.165 Μ (0.33 M carbon).

Cells were harvested in mid-exponential phase and incubated with lysozyme (20 mg/ml) for 2h at 25 °C. Total RNA isolation was performed using NucleoSpin RNA kit (Macherey-Nagel, Lab Supplies Scientific SA, Hellas) according to manufacturer guidelines. RNA samples were treated with DNase I as part of the kit procedure to eliminate any genomic DNA contamination. RNA concentration and purity were determined at 260 and 280 nm using an μDrop Plate with a Multiskan GO Microplate Spectrophotometer (Thermo Fisher Scientific, Waltham, MA USA), while RNA integrity was evaluated by agarose gel electrophoresis.

### First-strand cDNA synthesis

Reverse transcription took place in a 20 μL reaction containing 500 ng total RNA template, 0.5 mM dNTPs mix, 200U SuperScript II Reverse Transcriptase (Invitrogen, Antisel SA, Hellas), 40U RNaseOUT Recombinant Ribonuclease Inhibitor (Invitrogen, Antisel SA, Hellas) and 4 μM random hexamer primers (Takara Bio, Lab Supplies Scientific SA, Hellas). Reverse transcription was performed at 42 °C for 50 min, followed by enzyme inactivation at 70 °C for 15 min. The concentration of cDNA was determined using an μDrop Plate with a Multiskan GO Microplate Spectrophotometer (Thermo Fisher Scientific, Waltham, MA USA).

### Quantitative Real-Time PCR (qPCR)

qPCR assays were performed on the 7500 Real-Time PCR System (Applied Biosystems, Carlsbad, CA) using SYBR Green I dye for the quantification of *dszA, dszB, dszC, cbs,* and *metB* transcript levels. Specific primers were designed based on the published sequences of IGTS8 desulfurization operon (GenBank: U08850.1 for *dszABC*) and IGTS8 chromosome (GenBank: CP029297.1 for *cbs*, *metB, gyrB*) and are listed in Table S2. The gene-specific amplicons generated were 143 bp for *dszA*, 129 bp for *dszB*, 152 bp for *dszC*, 226 bp for *cbs,* 129 bp for *metB* and 158 bp for *gyrB*. The 10 μL reaction mixture included 5 μL Kapa SYBR Fast Universal 2x qPCR master mix (Kapa Biosystems, Lab Supplies Scientific SA, Hellas), 5 ng of cDNA template, and 200 nM of each specific primer. The thermal protocol was initiated at 95 °C for 3 min for polymerase activation, followed by 40 cycles of denaturation at 95 °C for 15 sec, and primer annealing and extension at 60 °C for 1 min. Following amplification, melt curve analyses were carried out to distinguish specific amplicons from non-specific products and/or primer dimers. All qPCR reactions were performed using two technical replications for each tested sample and target, and the average CT of each duplicate was used in quantification analyses, according to the 2^-ΔCCT^ relative quantification (RQ) method. The DNA gyrase subunit B (*gyrB*) gene from strain IGTS8 was used as an internal reference control for normalization purposes. A biological replicate of a cDNA sample derived from *R. qingshengii* IGTS8 grown on 1 mM DMSO for 66 h was used as our assay calibrator.

## Acknowledgments

We thank Jacob Bobonis (EMBL Heidelberg) for the S17-1 strain. This research project was supported by the Action RESEARCH – CREATE – INNOVATE co-financed by the European Regional Development Fund of the European Union and national resources through the Operational Program “Competitiveness, Entrepreneurship & Innovation” (EPAnEK) - NSRF (2014–2020) (Project code: T1EDK-02074, MIS 5030227).

## Supplementary Material

**Figure S1.**
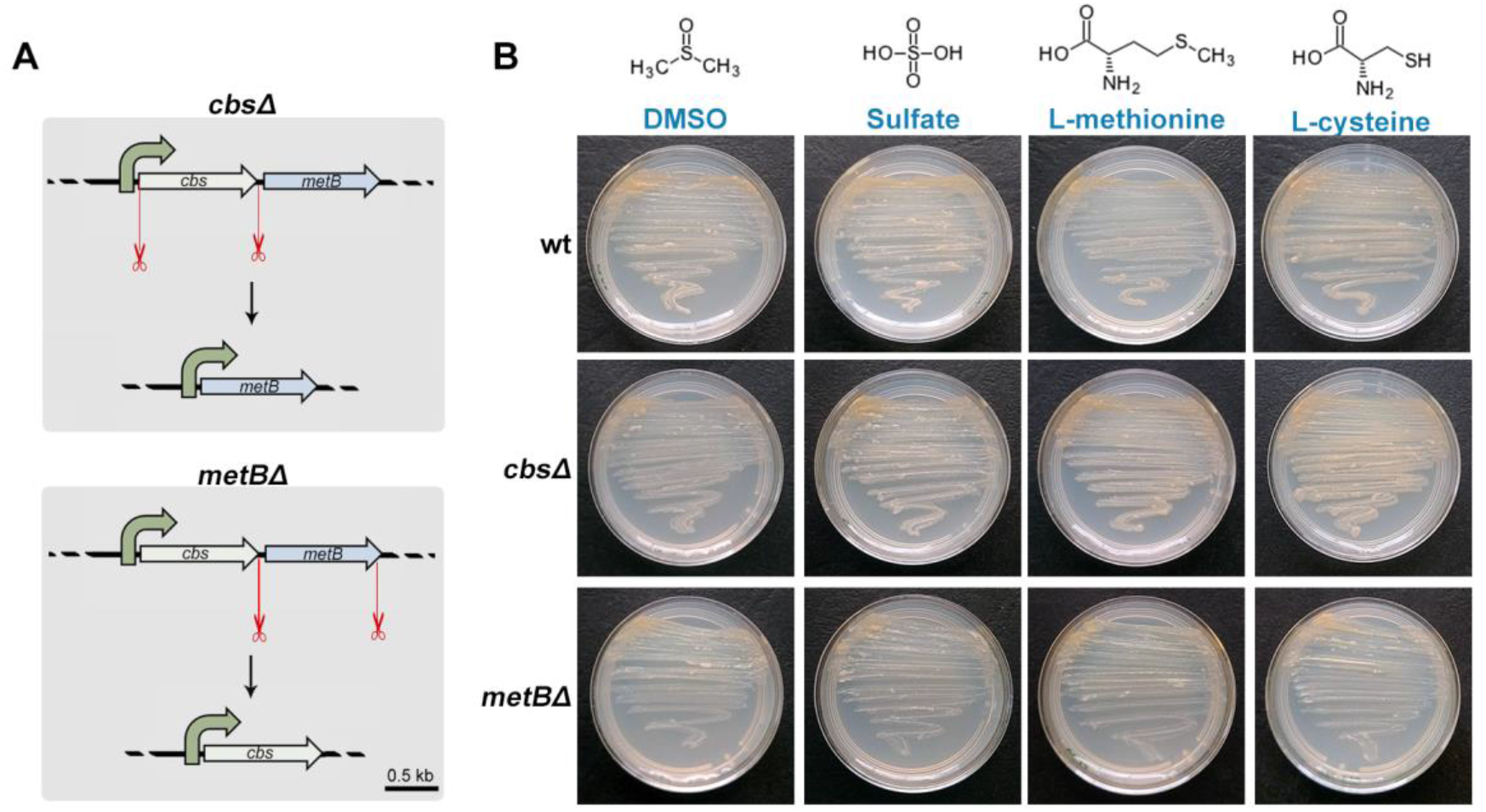
Cartoon and growth tests of strains *cbsΔ* and *metBΔ.* (A) Diagram of gene knockouts. Upper panel: *Targeted cbs* ORF deletion of 1386 bp. *metB* is expressed from the promoter sequence located in the upstream flanking sequence of the genetic locus. Lower panel: Deletion of 1168 bp within the *metB* ORF. (B) Growth tests of wild-type (wt) *R. qingshengii* IGTS8 and knockout strains *cbsΔ, metBΔ.* Basal minimal medium was supplemented with ethanol as a carbon source and 1mM of each sulfur source.

**Table S2.**
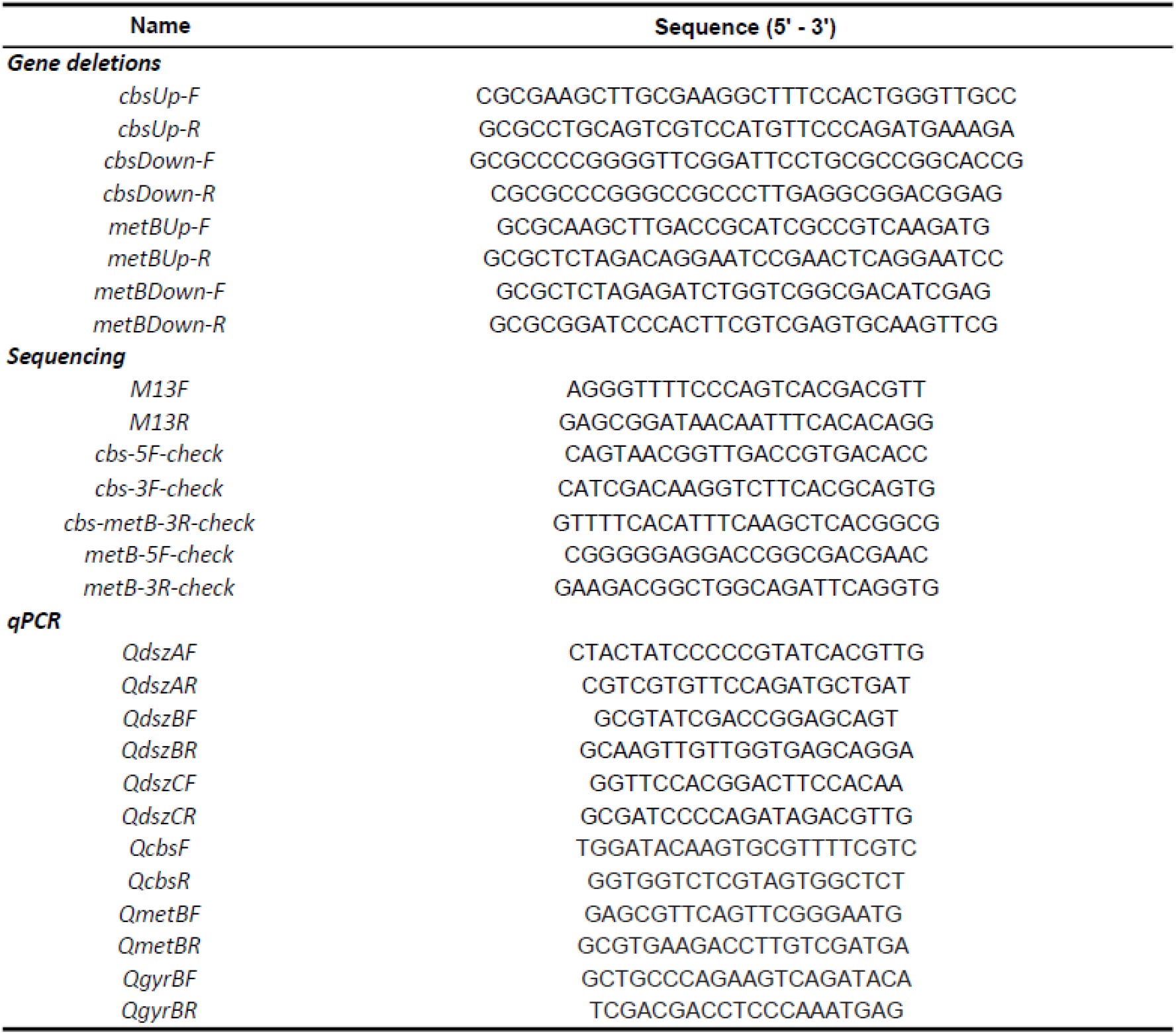
Oligonucleotides used in this study.

